# Conformational plasticity and dynamic interactions of the N-terminal domain of the chemokine receptor CXCR1

**DOI:** 10.1101/2020.12.17.423199

**Authors:** Shalmali Kharche, Manali Joshi, Amitabha Chattopadhyay, Durba Sengupta

## Abstract

Dynamic interactions between G protein-coupled receptors (GPCRs) and their cognate protein partners at the membrane interface control several cellular signaling pathways. An important example is the association of CXC chemokine receptor 1 (CXCR1) with its cognate chemokine, interleukin-8 (IL8 or CXCL8) that regulates neutrophil-mediated immune responses. Although the N-terminal domain of the receptor is known to confer ligand selectivity, the conformational dynamics of this intrinsically disordered region of CXCR1 in particular, and chemokine receptors in general, remains unresolved. In this work, we have explored the interaction of CXCR1 with IL8 by microsecond time scale coarse-grain simulations that were validated by atomistic models and NMR chemical shift predictions. We show that the conformational plasticity of the *apo-receptor* N-terminal region is restricted upon ligand binding, driving it to an open C-shaped conformation. Importantly, we validated the dynamic complex sampled in our simulations against chemical shift perturbations reported by previous NMR studies. Our results indicate that caution should be exercised when chemical shift perturbation is used as a reporter of residue contacts in such dynamic associations. We believe our results represent a step forward in devising a strategy to understand intrinsically disordered regions in GPCRs and how they acquire functionally important conformational ensembles in dynamic protein-protein interfaces.

**Author summary:** How cells communicate with the outside environment is intricately controlled and regulated by a large family of receptors on the cell membrane (G protein-coupled receptors or GPCRs) that respond to external signals (termed ligands). Chemokine receptors belong to this GPCR family and regulate immune responses. We analyze here the first step of binding of a representative chemokine receptor (CXCR1) with its natural ligand, interleukin 8 (IL8) by an extensive set of molecular dynamics simulations. Our work complements previous mutational and NMR experiments which lack molecular-level resolution. We show that in the inactive state, one of the extracellular domains of the CXCR1 receptor, namely the N-terminal domain, is highly flexible and like a “shape-shifter” can exist in multiple conformational states. However, when IL8 binds, the N-terminal domain undergoes a conformational freezing, and acquires a C-shaped “claw-like” structure. The complex between the receptor and IL8 is still quite dynamic as this C-shaped N-terminal domain forms an extensive but slippery interface with the ligand. We further validated these results by quantitative comparison with NMR and mutagenesis studies. Our work helps clarify the inherent disorder in N-terminal domains of chemokine receptors and demonstrates how this domain can acquire functionally important conformational states in dynamic protein-protein interfaces.

## Introduction

G protein-coupled receptors (GPCRs) are an important class of membrane-embedded receptors that respond to a diverse range of stimuli.^1,2^ These receptors play a central role in several cellular signaling pathways, and consequently are targeted by a large number of drugs.^3,4^ Recent advances in GPCR structural biology have helped to resolve the structure of transmembrane domains of several GPCRs. However, the interconnecting loops and the N-and C-terminal extramembranous regions remain largely unresolved.^5,6^ The high flexibility associated with these domains confers an intrinsic challenge in resolving specific conformational states of GPCRs, but attaches a functional significance to it.^6,7^ Both direct interaction *(e.g.,* between intracellular loop 3 (ICL3) and effectors^8^) and allosteric modulation by extramembranous loops (such as extracellular loops 2 and 3 (ECL2, ECL3))^6,9,10^ have been reported in various GPCRs. The N-terminal region, known to interact with ligands^11^ in GPCRs such as chemokine receptors,^12–14^ is of special interest in this context. In addition, N-terminal population variants of several GPCRs have been reported to alter drug response by allosteric modulation of ligand binding.^15–17^ Interestingly, lipid specificity and conformational sensitivity of extramembranous regions in GPCRs have recently been reported.^18–20^ In spite of their functional role, extramembranous regions in GPCRs remain largely uncharacterized in terms of their structure and dynamics.

Chemokine receptors are members of the GPCR superfamily that bind chemokine secretory proteins and play a fundamental role in innate immunity and host defense.^21,22^ These receptors highlight the functional importance of the N-terminal region since it represents the first site of ligand binding and confers selectivity to these receptors.^23^ A common two-site/two-step model has been proposed for chemokine binding that suggests interactions between receptor N-terminal domain and chemokine core (site-I) and between the chemokine N-terminus and receptor extracellular regions or transmembrane residues (site-II).^23–25^ In addition, recent reports confirm that the stoichiometry of binding is 1:1, although both the receptor and chemokines have been shown to dimerize in the cellular milieu.^24–26^ Early attempts to structurally characterize these complexes focused on site-I interactions and solution NMR approaches were successful in resolving the interactions between chemokines and short receptor fragments without the context of the full-length receptor or membrane environment.^27,28^ More recently, crystal structures have resolved site-II interactions, but only a partial site-I engagement.^29,30^ However, a superposition of structures with respect to the bound chemokine indicates that the placement of the receptor N-terminus could be receptor-specific.^31^ Although the two-site model served as the initial framework of functionally relevant interactions leading to chemokine-receptor binding, growing literature suggests a need for more complex models accounting for the dynamic mechanism of receptor-ligand binding.^32^

The CXC chemokine receptor-1 (CXCR1) is a representative chemokine receptor that controls the migration of neutrophils to infected tissues.^33^ The three-dimensional structure of CXCR1 (residues 29-324) has been elucidated by solid state NMR^34^ and follows a typical GPCR fold, with seven transmembrane α-helices interconnected by three intracellular and three extracellular loops. The two flanking domains, the extracellular N-terminal and intracellular C-terminal regions, were not resolved in this structure. CXCR1 binds the CXC ligand, CXCL8, commonly termed interleukin-8 (IL8). There are several reported structures of IL8 in monomeric and dimeric forms, although none bound to CXCR1.^27,28,35^ Several studies have highlighted a crucial role of the N-terminal region of CXCR1 in ligand binding affinity and selectivity.^36^ The interactions of IL8 were assessed using NMR with CXCR1 constructs of varying length, clearly indicating that IL8 could not bind to CXCR1 when the receptor N-terminal was truncated.^37^ In addition, IL8 was shown to bind with higher affinity to the CXCR1 N-terminal region in a lipid environment relative to that in solution,^36^ in agreement with our previous work using fluorescence and molecular dynamics (MD) simulations which show membrane interaction of the CXCR1 N-terminal region.^38–40^

In this work, we have examined chemokine-receptor interaction focusing on the N-terminal region of CXCR1 and its role in chemokine binding. We performed simulations of *apo*-CXCR1 as well as CXCR1 coupled with IL8 at coarse-grain and atomistic resolutions to monitor differential dynamics of the N-terminal region. We show that the N-terminal region is the first site of chemokine binding which restricts its conformational dynamics. The receptor-chemokine (CXCR1-IL8) complex consists of an extensive dynamic interface and we map the interactions both within the receptor and with the ligand. These results were further validated by comparison with chemical shift calculations reported in earlier NMR studies. Our results offer molecular insight into the interactions between CXCR1 and IL8, and would be useful in gaining a fundamental understanding of the initial events in chemokine-receptor interactions at site-I.

## Results

The N-terminal region of the chemokine receptor CXCR1 remains structurally unresolved in experiments due to its inherent flexibility.^34^ The importance of this region is reflected in reports that implicate it in the binding of the cognate chemokine (IL8),^36,37^ similar to all members of the chemokine receptor family.^14^ To explore the underlying molecular interactions, we have performed coarse-grain molecular dynamics simulations of CXCR1 and validated them against atomistic models. We report here the functional dynamics of the N-terminal region of CXCR1 in the *apo-* and IL8-bound forms.

### Conformational plasticity of the N-terminal region of *apo*-CXCR1

Coarse-grain simulations of the *apo*-CXCR1 receptor were performed starting from the extended N-terminal conformer (Fig 1a). In total, twenty simulations were performed totaling to 200 μs. During the simulations, the N-terminal region relaxed quickly from the initial structure and appeared more dynamic than the rest of the receptor. The N-terminal region sampled several orientations and was found to interact at different time points with the membrane bilayer and the transmembrane domains. The two main conformers observed (membrane-bound and receptor-contacted conformers) are shown in Figs 1b and 1c. These can be distinguished by the distance of distal residues 1-10 of the N-terminal region from the membrane (see Fig 1d). Several close interactions with the membrane (blue stretches) and multiple association-dissociation events were observed (see Fig 1d). When the N-terminal region dissociated from the membrane, it was located on top of the receptor, interacting with the transmembrane helices. In this state, it adopted a more compact conformation, as reflected in the radius of gyration (see Fig 1e). Overall, the position of the N-terminal region in the *apo*-receptor was highly dynamic.

**Fig 1.**
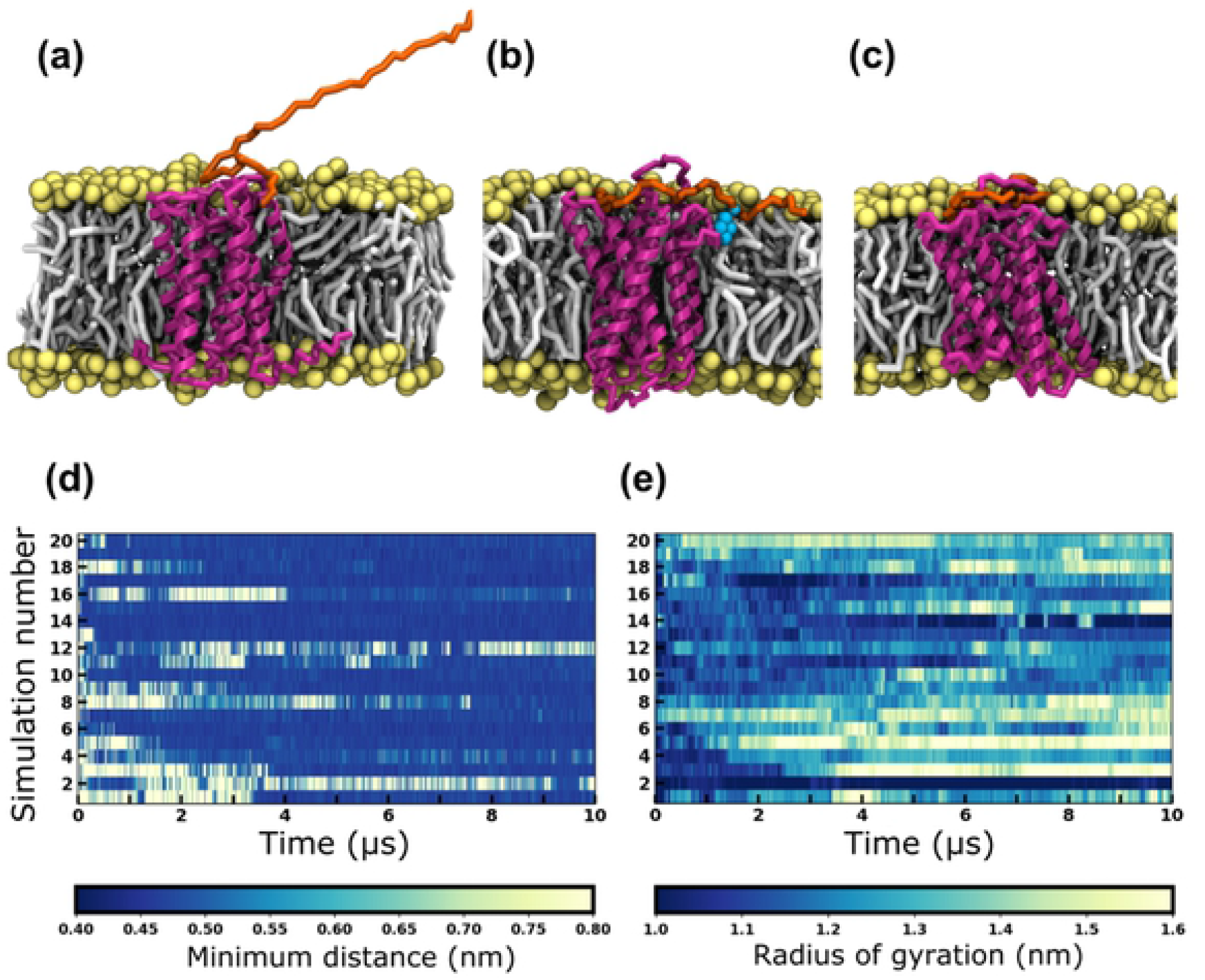
Representative snapshots of CXCR1 embedded in a lipid bilayer and membrane interaction of its N-terminal region. A visual representation of (a) the starting conformation with an extended N-terminal region, (b) the membrane-embedded N-terminal conformer and (c) the receptor-contacted N-terminal conformer. The receptor is depicted in magenta, the N-terminal region in orange, and the lipid headgroups and tails in yellow and gray, respectively. Water and ions are not displayed for clarity. The residue W10 of the N-terminal region, which interacts with the lipid bilayer is shown as cyan colored beads. (d) The minimum distance between the lipid bilayer and the distal part of the N-terminal region (residues 1-10) is plotted for 20 simulations of *apo*-CXCR1 as a function of time. The color bar denotes minimum distance in nm. A distance of ~0.4 nm (dark blue patches) indicates the binding of the N-terminal region to the lipid bilayer. (e) The radius of gyration of the N-terminal region is plotted for *apo*-CXCR1 as a function of time. The color bar denotes radius of gyration in nm. See Methods for more details.

To test the conformational landscape sampled in the coarse-grain simulations, we performed all-atom simulations of CXCR1 embedded in the membrane bilayer (see S1 Fig). The N-terminal region of CXCR1 adopted multiple conformations, and no stable secondary structure was observed over time (S1b Fig.). For a direct comparison, the intra-protein contacts were computed from both coarse-grain and atomistic simulations. Several off-diagonal elements were observed in both cases representing close interactions between residues which are sequentially apart (S1a Fig.). The off-diagonal contacts in the middle of the N-terminal region (around residues 20-25) indicate a compact conformation. Interestingly, we observed similar patterns in the contact maps (S1 Fig), indicating that the coarse-grain simulations were able to capture the overall conformational dynamics of this highly flexible region.

### The N-terminal region is the first site of ligand binding

We carried out coarse-grain simulations of CXCR1 with IL8 to examine the effect of ligand binding upon the structural dynamics of the N-terminal region of CXCR1. Overall, forty simulations were performed with two conformations of CXCR1 N-terminal region (membrane-bound and receptor-contacted) and two placements of IL8 (N-domain of the ligand facing the receptor and away from it). During the course of the simulations, IL8 diffused randomly in water and was observed to bind to the membrane-embedded CXCR1 within microseconds. A representative snapshot of the CXCR1-IL8 complex is shown in Fig 2a. The binding events were quantified from the minimum distance between IL8 and the receptor (Fig 2b and S2 Fig). The distance around 0.5 nm (blue stretches in the plot) indicate close interactions between the two proteins. A few binding-unbinding events were observed before the final bound complex was formed and no further unbinding was observed during the course of the simulations.

**Fig 2.**
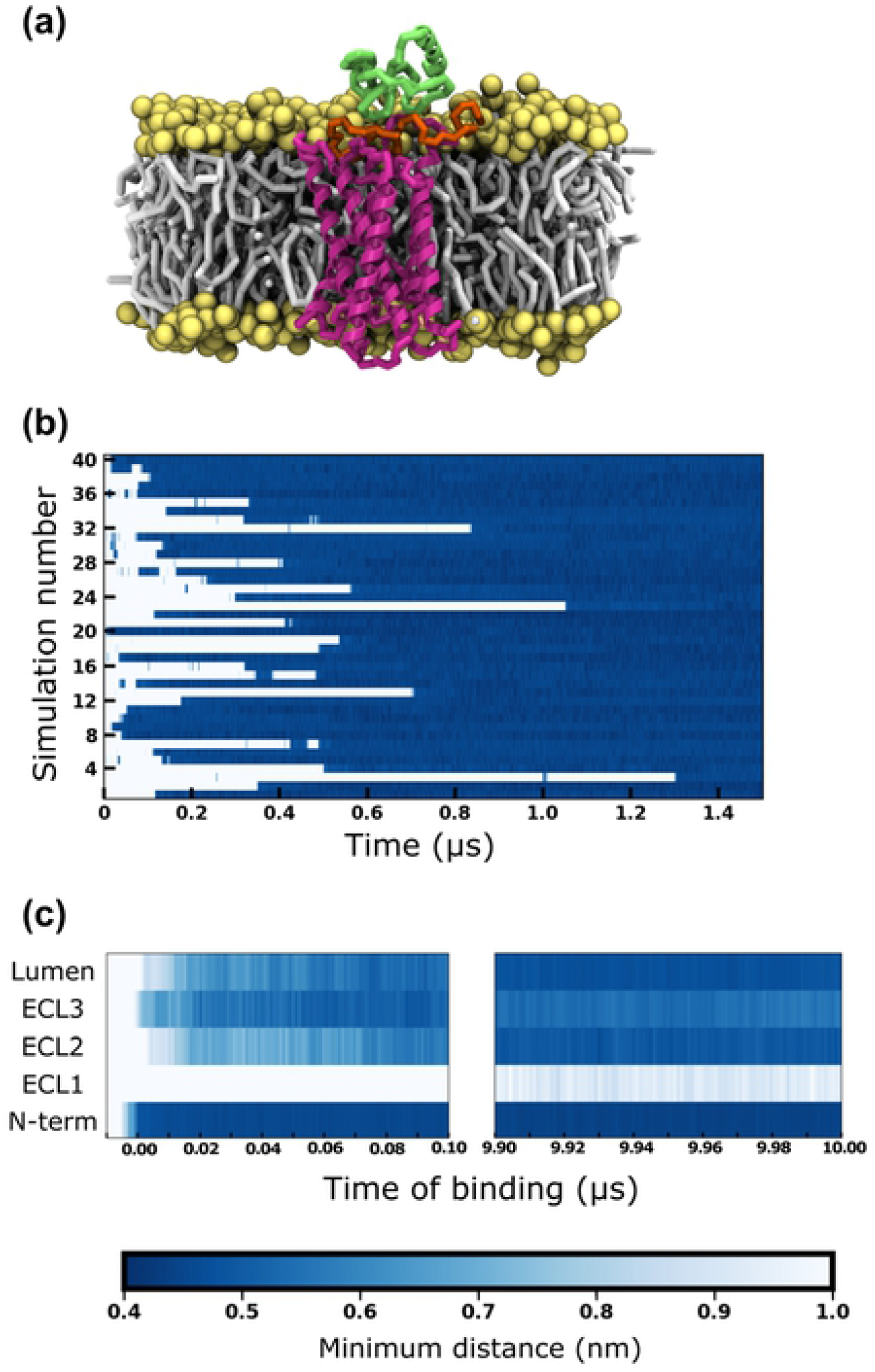
Interactions between the extracellular domains of CXCR1 and IL8. (a) A representative snapshot of IL8 bound to CXCR1. The receptor is shown in magenta, IL8 in green, and lipid headgroups and tails in yellow and gray, respectively. The N-terminal region of the receptor is highlighted in orange. Water molecules and ions are not shown for clarity. (b) The minimum distance (closest approach) between IL8 and CXCR1 plotted for the first 1.5 μs in forty simulations. The white stretches represent the unbound regime and the blue stretches represent the ligand-bound regime. Time of binding (t = 0) is defined as the time of first contact in the binding regime (0.5 nm distance cutoff) which remains undissociated till the end of the simulation. (c) The minimum distance between IL8 and various domains of the receptor as a function of time, considering the time of binding as t = 0. The values are averaged over all sets from the time of binding and plotted for the first 100 ns (left panel) and the last 100 ns (right panel). The color bar denotes minimum distance between IL8 and CXCR1 domains. See Methods for more details.

To understand the mechanism of binding, we characterized the interaction between the receptor domains and IL8 from the time of binding (Fig 2c). The time point corresponding to the binding event (time of binding t=0) is considered to be the time frame where the final bound complex is formed (taken from Fig 2b). For clarity, the receptor domains considered were the N-terminal region, the three extracellular loops (ECL1-3) and the lumen defined as the residues from the transmembrane helices lining the top of the receptor lumen. The minimum distance (distance of closest contact) between these domains and IL8 was calculated from the time of binding and averaged over all simulations. Interestingly, the N-terminal region was observed to be the first site involved in binding of IL8 (Fig 2c). Subsequently, IL8 was observed to interact with ECL3 followed by ECL2 and the lumen, and ECL1 does not appear to make any contacts. These contacts are maintained till the end of the simulations (10 μs after the initial binding) and the interactions with the N-terminal region appear quite stable. No interactions were observed with ECL1 consistent with the initial binding mode. We observed that the interactions with ECL3 reduced and that with ECL2 and the top of the lumen increased with time. We were unable to discern a deeper binding of the N-domain of IL8 in the receptor lumen. Overall, we observed that the N-terminal region of CXCR1 is the first site of binding for IL8 and this contact is maintained throughout the course of the simulations along with additional contact with sites on ECL2, ECL3 and the lumen.

### Conformational restriction in the N-terminal region upon ligand binding

To analyze the effect of ligand binding on conformational dynamics of the N-terminal domain, we computed intra-protein contact maps of the N-terminal region in the ligand-bound complex. These contact maps represent pair-wise probabilities of interaction for each residue pair within the N-terminal region, averaged over simulation time and all simulation sets. A composite contact probability map displaying direct comparison of residue-wise contacts within the N-terminal region from *apo*-CXCR1 (upper diagonal) and CXCR1-IL8 complex (lower diagonal) is shown in Fig 3. Interestingly, several intra-protein contacts observed in the *apo*-receptor appear to be lost in the ligand-receptor complex and the N-terminal region appears to be more open in the ligand-receptor complex. A few intra-protein contacts were observed in the distal region of the N-terminal region in the ligand-bound complex, but appear to be relatively weak. We identified six representative inter-residue contacts that dynamically form in the *apo*-receptor, but are completely absent in the ligand-bound simulations (S3 Fig). These interactions include electrostatic interaction (Met1-Asp26), putative hydrogen bonding (Thr5-Thr18, Ser2-Thr18) and aromatic ring stacking (Phe17-Tyr27).

**Fig 3.**
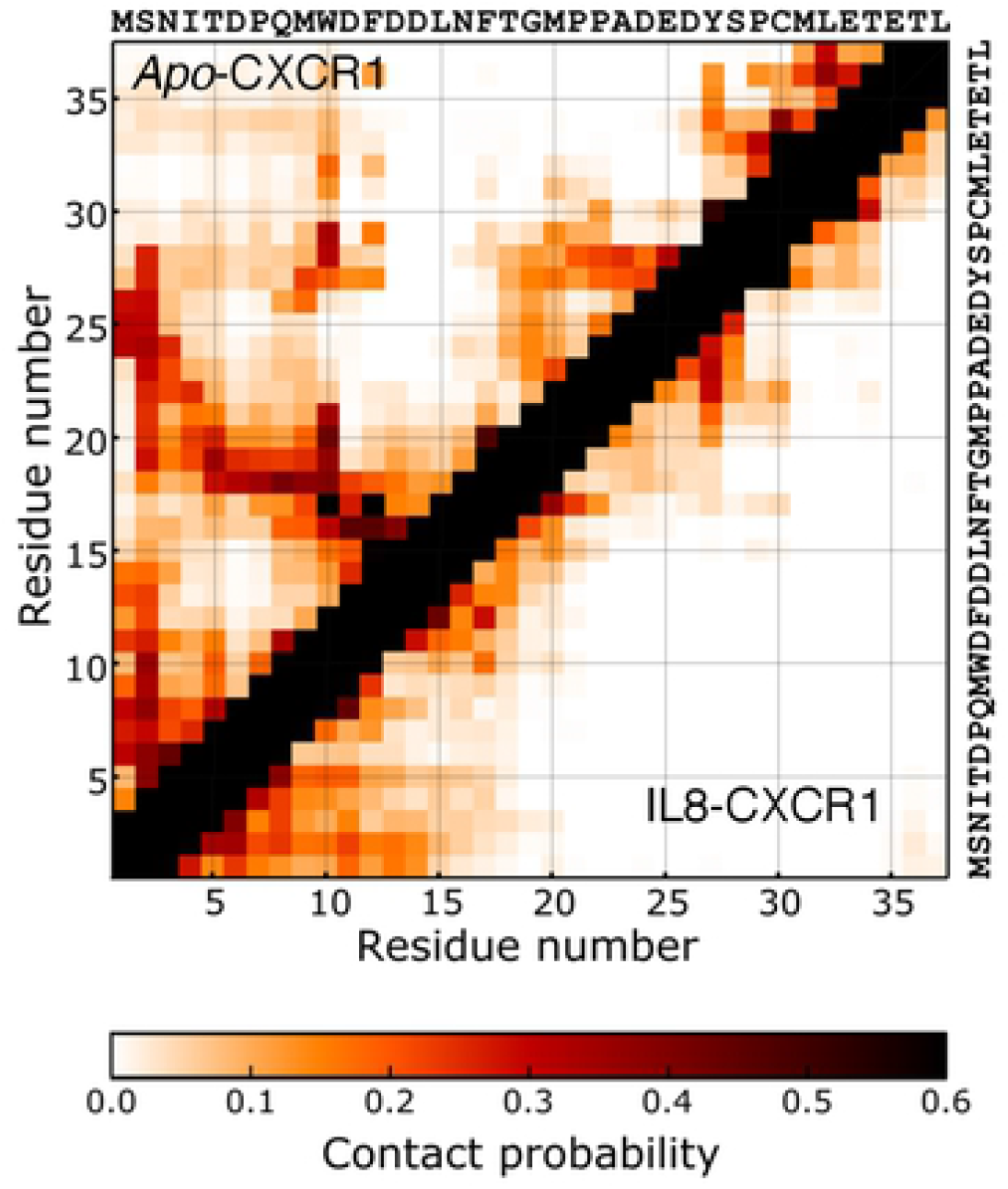
Conformational dynamics of the N-terminal region of CXCR1. Intra-protein contact maps of the N-terminal region of CXCR1 in presence (lower matrix) and absence (upper matrix) of the ligand. Residue-wise contact probabilities of the N-terminal region in *apo-* and IL8-bound CXCR1 are plotted in the top and bottom diagonal of the matrix, respectively. The amino acid sequence of the N-terminal region is displayed on the top and right. The values of contact probabilities (0.5 nm distance cutoff) are denoted in the color bar. See Methods for more details.

A more detailed characterization of the conformational dynamics was carried out by projecting the simulation trajectories onto a two-dimensional phase space. The two collective variables considered for the projection were the backbone RMSD of the N-terminal region and the distance distribution of an inter-residue contact Met1-Asp26 (Fig 4). The backbone RMSD describes an overall structural deviation with respect to a reference structure corresponding to the highest population cluster. The second reaction coordinate, *i.e.,* the distance between N-terminal residues Met1 and Asp26, reports on the end-to-end distance of the N-terminal region. Fig 4 shows the relative populations of the N-terminal region along these reaction coordinates sampled in the *apo*- and IL8-bound CXCR1 simulations. Multiple clusters were observed in the *apo*-receptor (marked I-III in Fig 4a), but only a single broad cluster (I) was observed in the ligand-bound receptor. The major cluster (cluster I in Fig 4b) in the IL8 bound simulations consists of conformers with a high end-to-end distance but low RMSD. The main cluster (cluster II in Fig 4a) in the *apo*-receptor exhibits a high RMSD. Interestingly, cluster I in the *apo*-receptor appears to overlap with a part of the conformational space sampled in the ligand-bound complex.

**Fig 4.**
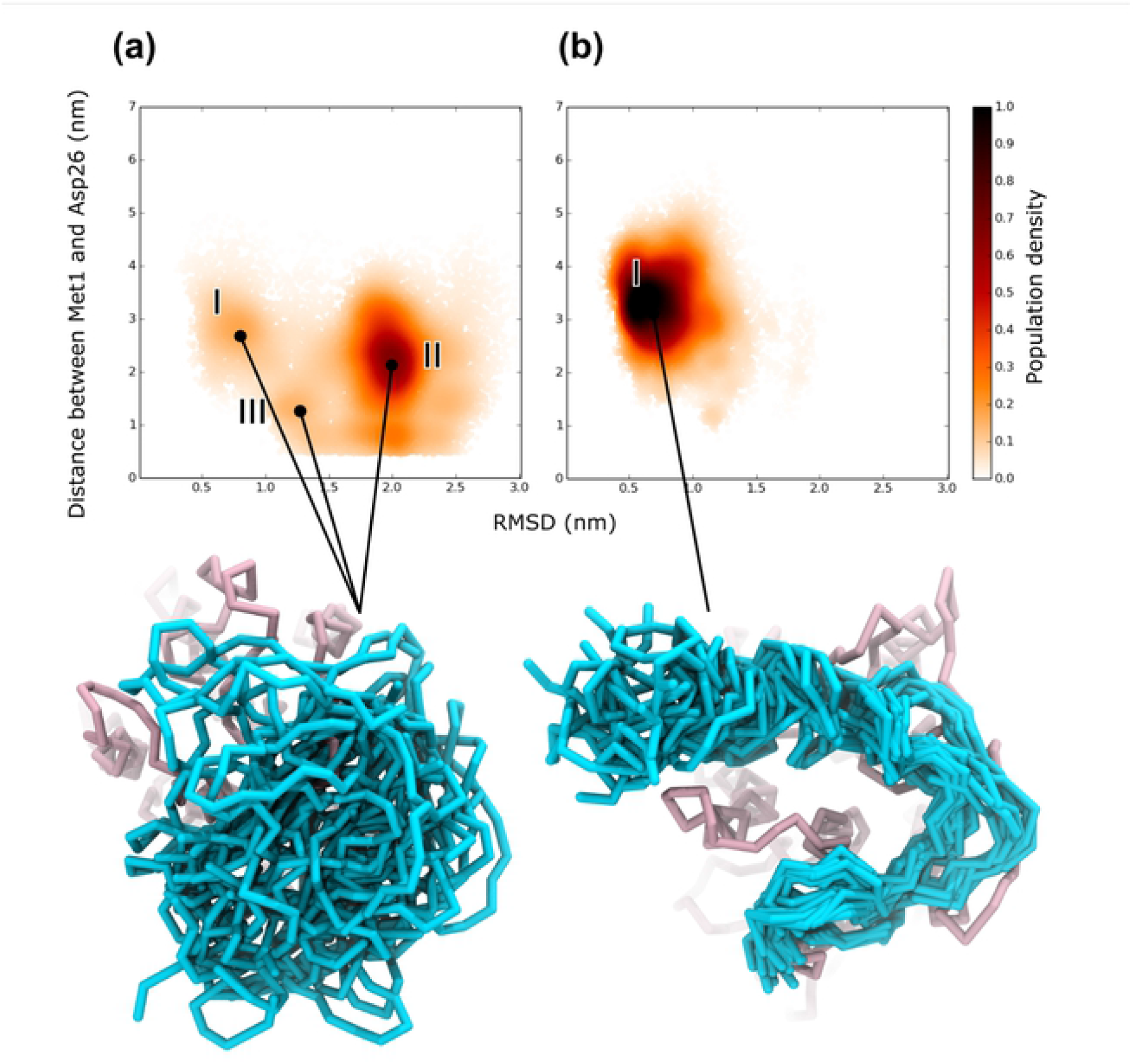
Conformational landscape of the N-terminal region of CXCR1. Population density map of the conformations sampled by the N-terminal region plotted as a function of backbone RMSD of the N-terminal region and the distance between side chains of two representative residues (Met1 and Asp26) for (a) *apo*-CXCR1 and (b) IL8-bound CXCR1. The most populated conformations are shown below the plots. The N-terminal region is shown in cyan and rest of the receptor is in pink. See Methods for more details.

The single cluster in the ligand-bound complex (Fig 4b) appears to be in contrast to the lack of intra-protein contacts observed in the receptor-ligand simulations (see Fig 3). A visual inspection revealed that the ligand-bound structures adopt a C-shape in the N-terminal region (Fig 4b). Such a conformation allows a more extensive protein-protein interface when the ligand is bound to the receptor, but at the same time results in the loss of intra-protein contacts. To characterize this C-shaped state, we calculated the contact maps of the interactions between the N-terminal region and the extracellular loops (S4 Fig). Interestingly, we observed large differences in the interactions in the *apo-* and IL8-bound N-terminal region. The N-terminal region of the *apo*-receptor samples several interaction sites on the extracellular loops and we could not discern a consensus pattern of interacting residues, confirming the presence of diverse conformational states. In contrast, specific regions of the N-terminal region were found to interact with each of the extracellular loops in case of IL8-bound receptor, giving rise to a C-like shape.

### Mapping the N-terminal region interactions: Validation by chemical shift perturbations

We analyzed the molecular interactions of the N-terminal region by calculating the contact probabilities with the chemokine (see Fig 5a). We observed an extensive contact surface between the ligand and the N-terminal region, and a large number of flexible contacts were observed along the length of the N-terminal region. The contact map is consistent with the C-shaped N-terminal region described above with maximal contact probabilities at the center of the region. In particular, a high contact probability is observed at residues 20-25. The residues predicted to have a high contact probability match well with previous mutagenesis data. In particular, residues Pro21 and Tyr27 have been previously shown by mutational studies to be critical for ligand binding.^41^

**Fig 5.**
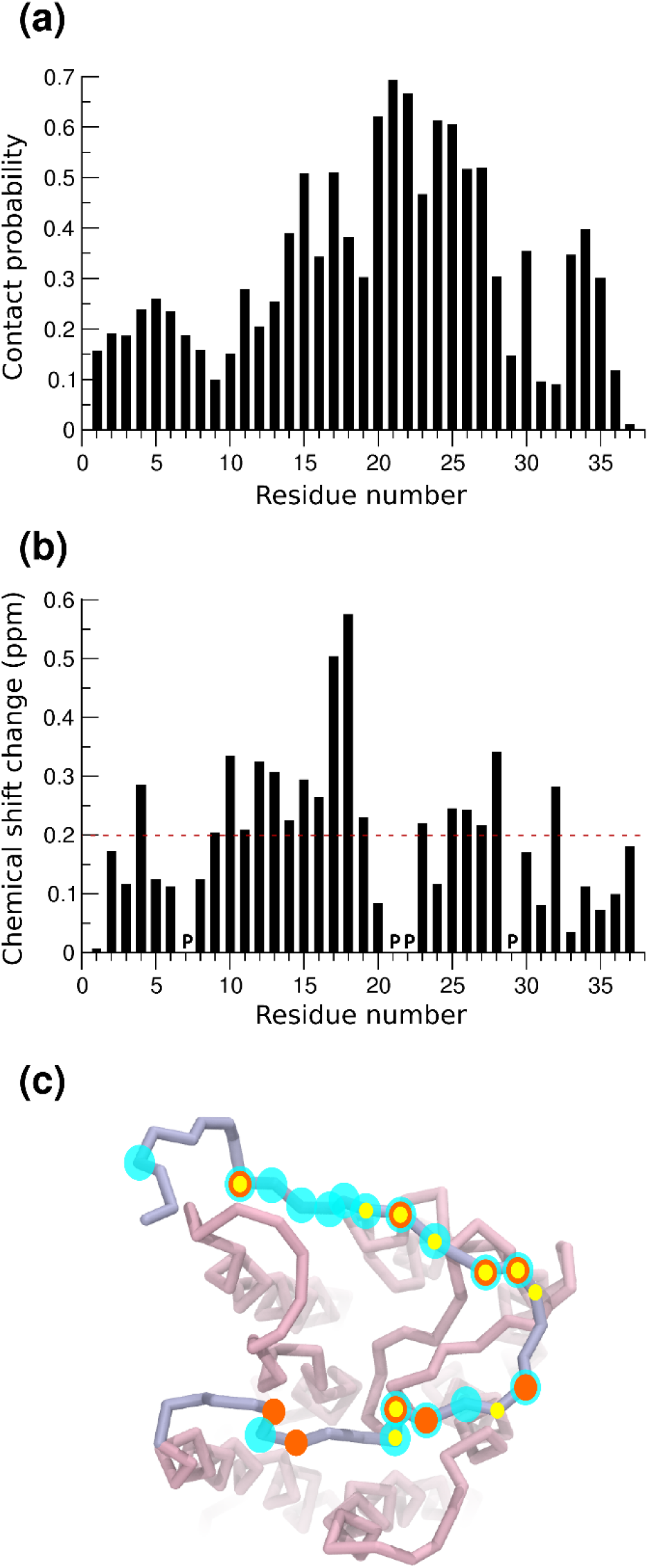
Residue-wise interactions of the N-terminal region of CXCR1 with IL8. (a) Residuewise contact probabilities of the N-terminal region interacting with IL8. (b) Predicted chemical shift changes in the N-terminal region between the *apo*- and ligand-bound state. (c) The N-terminal residues with chemical shift perturbations above a cutoff (dotted lines in panel (b)) mapped onto the receptor structure. The cyan transparent spheres represent residues from the predictions. The orange and yellow spheres represent residues showing significant chemical shift changes as reported from NMR measurements.^37,42^ See Methods and text for more details.

One of the few experimental approaches that are able to report conformational dynamics of this region is NMR using chemical shifts of the backbone amides that are closely related to their conformations. Chemical shift perturbations between the *apo*- and IL8-bound CXCR1 receptor from NMR studies in lipid environments have previously been reported.^37,42^ To compare this data with simulations reported here, we chose representative structures from each of the coarse-grain simulation sets and mapped them to their atomistic representation.

Subsequently, we computed the predicted chemical shifts in the backbone amides of N-terminal region using eq (1). The resultant chemical shift perturbations plotted as a function of residue number are shown in Fig 5b. We observe that the central segment of the N-terminal region (residues 10-19) shows a higher chemical shift perturbation. Residues at the distal and proximal end (residues 1-5 and 33-37) exhibit relatively lower perturbation. These perturbations arise both due to direct contacts with the ligand as well as conformational changes occurring in the N-terminal region upon ligand binding. Overall, we found a good agreement between the residues predicted in this work from simulations to have a large chemical shift perturbation and those reported earlier using NMR. These residues are pictorially depicted in Fig 5c. The residues highlighted in cyan were predicted by simulations to have a large chemical shift and residues in orange and yellow have been identified in previous experiments.^37,42^ We observe a considerable overlap in these residues, although many more residues were predicted to have a large chemical shift perturbation from our simulations relative to those identified using NMR. Nonetheless, a remarkable consistency is observed in the chemical shift perturbations predicted from coarse-grain simulations and those determined from NMR studies.

Interestingly, the chemical shift perturbations do not exactly match the interactions identified between the CXCR1 N-terminal region and the ligand from our simulations. In particular, a comparison of Figs 5a and 5b shows that residues 20-25 have a high contact probability, but low chemical shift perturbations. Similarly, residues 17-20 exhibit higher chemical shift difference relative to the corresponding contact probability. It is apparent that these chemical shift perturbations include environment effects due to altered conformational dynamics of the N-terminal region, particularly due to the C-shaped conformer adopted in the ligand-bound form. Since chemical shift perturbations are often used as a direct reporter of protein-protein contacts, we propose that caution should be exercised while interpreting such data, especially for intrinsically disordered regions. We believe that a combined approach integrating NMR and MD simulation approaches could provide novel insight into functional GPCR-ligand dynamics.

### Dynamic protein interactions define the chemokine N-domain and receptor interface

The dynamic interactions reflected in the contact probabilities at the CXCR1 N-terminal region (see Fig 5a) were observed in the ligand as well. We clustered the conformers corresponding to the different binding modes of IL8 with the CXCR1 N-terminal region. The five clusters that were observed to be most populated are shown schematically in Fig 6a. Overall, it appears that the receptor N-terminal wraps around the ligand (IL8) and explores several binding modes. The main binding mode (~40% population) indicates that maximal interactions are localized with the N-domain and α-helix of IL8. The second and third binding mode additionally involves β1 and β3 strands, respectively. Residues involved in maximal contact of IL8 with the N-terminal region of CXCR1 were identified and mapped onto the structure, along with residues reported from NMR^37,42^ and mutagenesis experiments^43–47^ (Fig 6b). As expected, residues from the N-domain and α-helix were found to be involved, together with residues from the β1 and β3 strands, in IL8-CXCR1 N-terminal domain interaction. Importantly, we found an overlap between the regions in IL8 predicted to interact with the receptor and those reported previously. However, the N-terminal residues predicted to be important from mutagenesis studies^43–47^ were not observed in our simulations or NMR studies^37,42^. The conformational plasticity of CXCR1 N-terminal region and dynamic interfaces sampled in the protein-protein complex appear to be a hallmark of chemokine-receptor binding.

**Fig 6.**
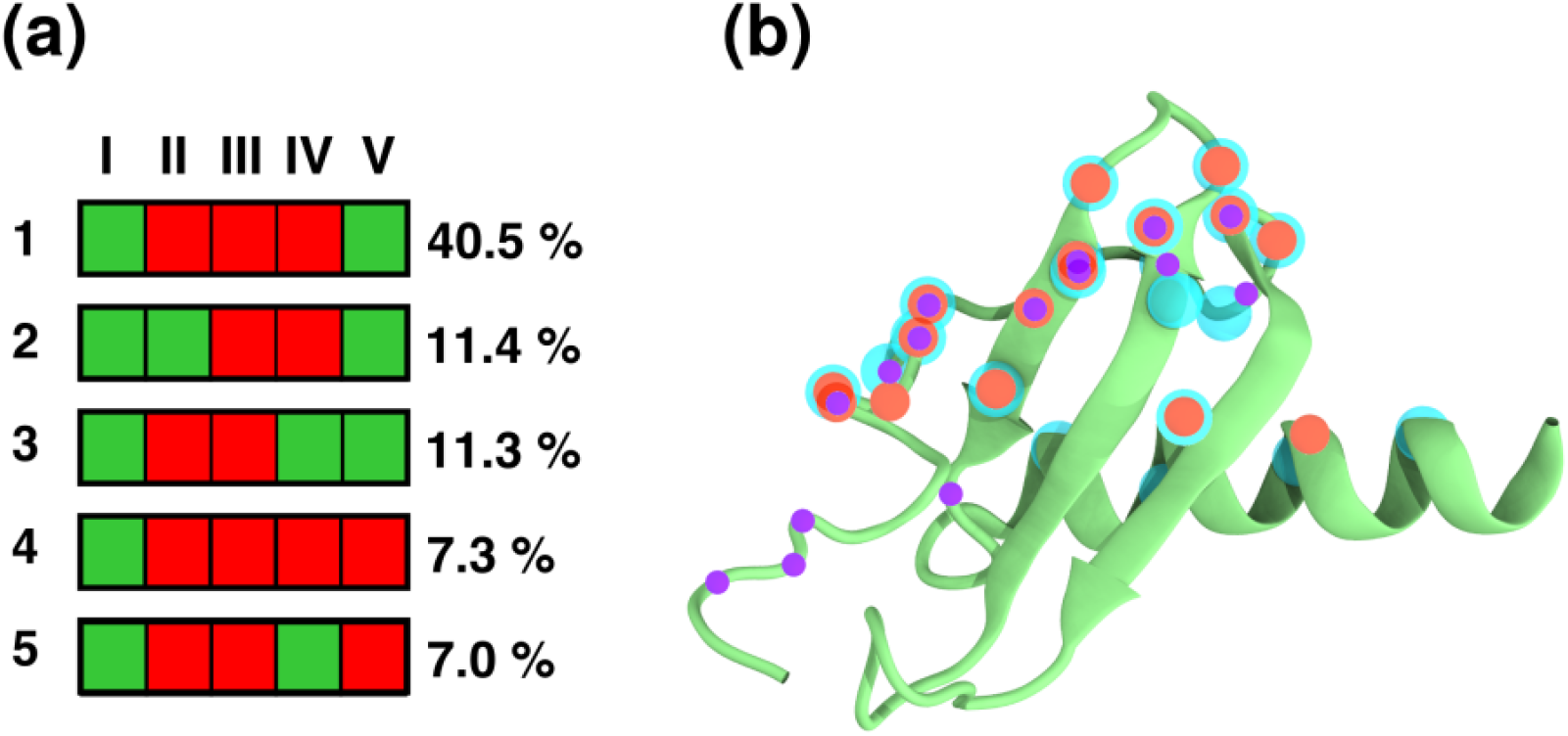
Binding modes of IL8 characterizing its interactions with the N-terminal region of CXCR1. (a) The most populated binding modes of IL8 characterized by the contacts formed by each of its structural element with the N-terminal region of CXCR1. The structural elements are denoted as I: N-domain, II: β1-strand, III: β2-strand, IV: β3-strand, and V: α-helix. The binding modes are numbered 1 to 5, in decreasing order of population. The green and red boxes represent interacting and non-interacting regions, respectively. (b) IL8 residues involved in binding to CXCR1 mapped on the cartoon representation of IL8. The cyan spheres represent interacting residues identified from our simulations. The orange and violet spheres represent interacting residues determined from previous NMR^37,42^ and mutagenesis^43–47^ studies, respectively. See Methods and text for more details.

## Discussion

The chemokine family of receptors are an important class of GPCRs that bind to the chemokine signaling proteins *via* their extracellular domains with a partial involvement of the transmembrane helices.^23^ A molecular resolution of CXCR1-IL8 interactions would open up avenues for therapeutic design and an overall understanding of immune signaling. In this work, we have addressed the molecular details underlying chemokine-receptor interactions focusing on the representative pair, CXCR1-IL8. In particular, we have analyzed the structural dynamics of the N-terminal region of CXCR1 in both *apo-* and ligand-bound forms. In the *apo*-receptor, the N-terminal region is highly dynamic, consistent with the absence of resolution by NMR^34^ and in agreement with its intrinsically disordered nature.^38,39^ Upon ligand binding, the N-terminus adopts a dynamic C-shaped conformation that facilitates ligand binding *via* an extensive and dynamic surface. Our results are in overall agreement with chemical shift differences reported from NMR studies. Taken together, our results represent an important step toward understanding chemokine-receptor interactions, especially with respect to the first site of binding.

An important finding from our work is the inherent conformational dynamics of the N-terminal region and the binding interface. The identification and prediction of molecular details underlying such protein-protein interfaces is challenging in the context of GPCR-ligand interactions. In mechanistic terms, the main challenges are (i) resolving distinct temporal/spatial interactions (two-site/two-step model), (ii) accounting for the dynamics of the intrinsically disordered N-terminal region, and (iii) inherent technical difficulties in resolving the structural dynamics of membrane receptors. We observed differential conformational dynamics sampled by the N-terminal region in the presence and absence of the ligand. Interestingly, the *apo*-receptor samples a sub-space overlapping with the IL8-bound N-terminal region dynamics (Fig. 4), suggesting a conformational selection by the ligand in the *apo*-receptor. Counterintuitively, the larger dynamics in the *apo*-receptor is associated with increased intra-protein contacts, whereas the C-shaped ligand-bound complex exhibits reduced intra-protein contacts. These loss of contacts within the N-terminal region in the IL8-bound complex are replaced by ligand contacts in the dynamic ligand-receptor interface. The dynamic protein-protein interface observed here represents an important aspect in the emerging understanding of plasticity in GPCR complexes.^48^

We observe that the N-terminal region is the first site of ligand binding in the CXCR1 receptor, consistent with models based on previous fluorescence and NMR studies.^36,37^ In the simulations, the chemokine adopts a peripheral arrangement and a deeper binding of N-domain in the receptor lumen was not observed. This mode of binding differs from crystal structures of other chemokine receptors, but is consistent with CXCR1 NMR data.^37^ In addition, a recent cryo-EM structure of a ternary complex of CXCR2, IL8 and G-protein reports that IL8 displayed a shallow binding mode compared to the other co-crystal structures of chemokines and their receptors.^49^-The extensive contact surface between the ligand and the receptor N-terminal region are consistent with recent hypothesis from experimental approaches in related receptors.^50^ In this work, we have compared chemical shift perturbations predicted from our simulations with results from NMR studies. Although the overall trends match quite well, we believe that the differences in the quantitative values could arise from the differential ensemble averages of experiments and simulations (due to different time scales associated with these approaches), peptide constructs used in experiments, and inaccuracies in prediction tools. In this context, we would like to recommend that caution should be exercised in assigning residues with high chemical shift perturbations to binding sites in receptors.^51^ Our results clearly show that the residues with maximum interactions do not necessarily exhibit the highest chemical shift perturbation. Instead, altered conformational dynamics of receptor N-terminal region (as reported here) could influence the observed chemical shift perturbations.

Computational studies, in close link with experimental approaches, have attempted to overcome some of the resolution problems associated with structure-based experiments. Several studies have combined docking followed by short MD simulations^52,53^ and have been able to capture important interactions, such as electrostatic interactions at site-I. Computational design of chemokine binding proteins, such as receptor-derived peptides capture agents from the extracellular domains of CXCR1^53^ has also been reported. Similar approaches combining docking with free energy calculations were used to design IL8-based peptide inhibitors to inhibit binding of CXCR1.^54^ To circumvent the problem of limited sampling, coarse-grain simulations coupled with replica exchange have been successfully used for predicting conformational ensembles associated with the binding of cyclic peptide antagonist to CXCR4.^55^ Coarse-grain simulations, in particular, appear to be well suited to predict proteinprotein interactions within the membrane, such as in single transmembrane helical receptors^56,57^ and GPCRs.^58–61^

In conclusion, we have used a combined atomistic and coarse-grain simulation approach to analyze the mechanism of binding of the chemokine IL8 to its cognate receptor CXCR1. We were able to observe the dynamic interfaces formed during the binding of CXCR1 and IL8. In addition, our results show that a conformational restriction of the flexible N-terminal region of the receptor induced by the ligand governs chemokine binding. These results suggest a conformational selection by the chemokine during the binding. The complementarity in shape and dynamic protein-protein interface appears to drive chemokine recognition by the receptor. We believe that our results represent an important step toward robust analysis of complex GPCR-ligand interactions and in designing improved therapeutics.

## Methods

### System setup and simulation parameters

The sequence of human CXCR1 N-terminal region (residues 1-37) was taken from the UniProtKB database (ID: P25024) and the structure was modeled in an extended conformation using Discovery Studio 3.5 (Accelrys Software Inc., Release 3.5, San Diego, CA). The *apo-* CXCR1 structure considered in this study was built by coupling the modeled structure of N-terminal domain to the NMR structure of CXCR1 (PDB ID 2LNL: residues 38-324). The energy of final atomistic structure was minimized (50,000 steps) using the steepest descent method. The structure was then mapped to its coarse-grain representation using parameters from the Martini v2.1 force field.^62,63^ The receptor was embedded in a pre-equilibrated 1-palmitoyl-2-oleoyl-*sn*-glycero-3-phosphocholine (POPC) bilayer (284 lipids) using insane.py script^64^ and then solvated. Twenty replicate simulations of 10 μs each were carried out for *apo*-CXCR1. The conformations of the N-terminal region sampled during these simulations were clustered, and two distinct receptor conformations were chosen, one with the N-terminal coiled on the top of the receptor (receptor-contacted) and other with the N-terminal interacting with the membrane bilayer (membrane-bound). For the ligand binding simulations of the two conformers (receptor-contacted and membrane-bound), IL8 was inserted at a distance of ~3 nm from the receptor to avoid potential bias arising from pre-placement. We considered two different orientations of IL8 while building these setups, resulting in four unique starting configurations of the CXCR1-IL8 simulations. The coarse-grain representation of IL8 was obtained by mapping from the atomistic three-dimensional structure (PDB ID: 1ILQ). Forty simulations of 10 μs each were run from these starting structures, both with and without elastic potential functions to fix the structural domains in IL8.^65^ The remaining parameters and setup were same as that of the CXCR1-IL8 system. The total simulation time was 400 μs, corresponding to 1.6 ms of atomistic sampling time.

All simulations were performed using the GROMACS-4.5.5 package.^66,67^ For coarse-grain simulations, Martini force field (versions 2.0 and 2.2)^62,63^ was used to represent lipids and proteins, respectively. Standard parameters corresponding to the coarse-grain Martini simulations were used. Non-bonded interactions were modeled using a cutoff of 1.2 nm. Electrostatic interactions were shifted to zero in the range 0 to 1.2, whereas Lennard-Jones potential was shifted to zero in the range of 0.9 to 1.2. Temperature was coupled to a thermostat at 300 K with a coupling constant of 0.1 ps using the v-rescale thermostat.^68^ Pressure was coupled at 1 bar with a coupling constant of 0.5 ps using the semi-isotropic Berendsen algorithm^69^ independently in the plane of the bilayer and perpendicular to the bilayer. Production runs were performed with a time step of 20 fs. Initial velocities for the systems were randomly chosen from a Maxwell distribution at 300 K.

The atomistic model of *apo*-CXCR1 was used as a starting structure for the all-atom MD simulations. The receptor was inserted in a pre-equilibrated POPC bilayer using the CHARMM-GUI module.^70^ Water and chloride ions were added to solvate and neutralize the charge on the system. Energy minimization was performed to remove steric clashes. The system was equilibrated under NVT conditions for 100 ps, followed by NPT ensemble for 1 ns, with position restraints on the receptor backbone. A production run of 1 μs was carried out as a control. In the atomistic simulations, temperature coupling was applied with the v-rescale thermostat^68^ to maintain temperature at 300 K. Semi-isotropic pressure coupling was applied to maintain a pressure of 1 bar along the direction of bilayer plane and perpendicular, using Parrinello-Rahman barostat.^71^-The long-range electrostatic interactions were treated with the particle mesh Ewald (PME) algorithm. The short-range electrostatic interactions and Lennard-Jones interactions were cutoff at 1.2 nm. A time step of 2 fs was considered for atomistic simulations.

### Analysis

Simulations were analyzed using in-house scripts, VMD^72^ and GROMACS utilities. The residue-wise contacts were calculated using the g_distMat tool *(https://github.com/rjdkmr/g_distMat).* For a given pair of residues, a contact was defined if the minimum distance between the residues (distance of closest approach) was within the cutoff (0.6 nm). The contact probability was calculated for each residue pair as the time for which they were in contact, normalized over the simulation length and averaged across all the simulation replicates.

To calculate chemical shift changes in the CXCR1 N-terminal region upon IL8 binding, we considered the main structures sampled in the coarse-grain simulations by clustering the conformations from each simulation replicate and a single conformer from each set was chosen. These conformers were transformed to the atomistic description (CHARMM36 force field) using Martini analysis tools.^64^ These structures were provided as an input to the SHIFTX2 program^73^ which predicts chemical shifts of backbone amides. The chemical shift values were averaged over replicates and chemical shift changes were calculated using the equation:

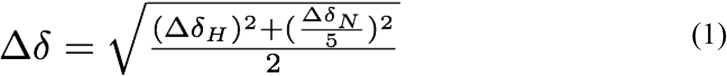

where Δδ_H_ is the change in the backbone amide proton chemical shift and Δδ_N_ is the change in the backbone amide nitrogen chemical shift.

## Acknowledgments

We gratefully acknowledge computing resources from CSIR-NCL and CSIR-Fourth Paradigm Institute. S.K. thanks the Council of Scientific and Industrial Research, Govt. of India, for the award of a Senior Research Fellowship. A.C. thanks Sreetama Pal for help and discussion during the preparation of the manuscript. We thank members of the Chattopadhyay laboratory for critically reading the manuscript and for their comments.

## Author Contributions

**Conceptualization:** Durba Sengupta, Manali Joshi, Amitabha Chattopadhyay.

**Formal analysis:** Durba Sengupta, Shalmali Kharche.

**Funding acquisition:** Durba Sengupta, Amitabha Chattopadhyay.

**Investigation:** Shalmali Kharche.

**Methodology:** Durba Sengupta, Shalmali Kharche.

**Resources:** Durba Sengupta, Amitabha Chattopadhyay.

**Supervision:** Durba Sengupta, Manali Joshi.

**Writing – original draft:** Shalmali Kharche, Durba Sengupta.

**Writing – review & editing:** Durba Sengupta, Amitabha Chattopadhyay.

## References

1. Rosenbaum DM, Rasmussen SGF, Kobilka BK. The structure and function of G-protein-coupled receptors. Nature. 2009;459: 356–363.

2. Pierce KL, Premont RT, Lefkowitz RJ. Seven-transmembrane receptors. Nat Rev Mol Cell Biol. 2002;3: 639–650.

3. Jacobson KA. New paradigms in GPCR drug discovery. Biochem Pharmacol. 2015; 98: 541–555.

4. Hauser AS, Attwood MM, Rask-Andersen M, Schiöth HB, Gloriam DE. Trends in GPCR drug discovery: new agents, targets and indications. Nat Rev Drug Discov. 2017;16: 829–842.

5. Venkatakrishnan AJ, Flock T, Prado, DE, Oates ME, Gough J, Babu MM. Structured and disordered facets of the GPCR fold. Curr Opin Struct Biol. 2014;27: 129–137.

6. Pal S, Chattopadhyay A. Extramembranous regions in G protein-coupled receptors: cinderella in receptor biology? J Membr Biol. 2019;252: 483–497.

7. Chattopadhyay A. GPCRs: lipid-dependent membrane receptors that act as drug targets. Adv Biol. 2014;2014: 143023.

8. Szczepek M, Beyrière F, Hofmann KP, Elgeti M, Kazmin R, Rose A, et al. Crystal structure of a common GPCR-binding interface for G protein and arrestin. Nat Commun. 2014;5: 4801.

9. Unal H, Karnik SS. Domain coupling in GPCRs: the engine for induced conformational changes. Trends Pharmacol Sci. 2012;33: 79–88.

10. Peeters MC, van Westen GJP, Li Q, IJzerman AP. Importance of the extracellular loops in G protein-coupled receptors for ligand recognition and receptor activation. Trends Pharmacol Sci. 2011;32: 35–42.

11. Coleman JLJ, Ngo T, Smith NJ. The G protein-coupled receptor N-terminus and receptor signalling: N-tering a new era. Cell Signal. 2017;33: 1–9.

12. Prado GN, Suetomi K, Shumate D, Maxwell C, Ravindran A, Rajarathnam K, et al. Chemokine signaling specificity: essential role for the N-terminal domain of chemokine receptors. Biochemistry. 2007;46: 8961–8968.

13. Ravindran A, Joseph PRB, Rajarathnam K. Structural basis for differential binding of the interleukin-8 monomer and dimer to the CXCR1 N-domain: role of coupled interactions and dynamics. Biochemistry. 2009;48: 8795–8805.

14. Szpakowska M, Fievez V, Arumugan K, van Nuland N, Schmit J-C, Chevigné A. Function, diversity and therapeutic potential of the N-terminal domain of human chemokine receptors. Biochem Pharmacol. 2012;84: 1366–1380.

15. Hauser AS, Chavali S, Masuho I, Jahn LJ, Martemyanov KA, Gloriam DE, et al. Pharmacogenomics of GPCR drug targets. Cell 2018;172: 41–54.e19.

16. Shahane G, Parsania C, Sengupta D, Joshi M. Molecular insights into the dynamics of pharmacogenetically important N-terminal variants of the human β2-adrenergic receptor. PLoS Comput Biol. 2014;10: e1004006.

17. Bhosale S, Nikte SV, Sengupta D, Joshi M. Differential dynamics underlying the Gln27Glu population variant of the β2-adrenergic receptor. J Membr Biol. 2019;252: 499–507.

18. Prasanna X, Jafurulla M, Sengupta D, Chattopadhyay A. The ganglioside GM1 interacts with the serotonin_1A_ receptor via the sphingolipid binding domain. Biochim Biophys Acta Biomembr. 2016;1858: 2818–2826.

19. Pal S, Aute R, Sarkar P, Bose S, Deshmukh MV, Chattopadhyay A. Constrained dynamics of the sole tryptophan in the third intracellular loop of the serotonin_1A_ receptor. Biophys Chem. 2018;240: 34–41.

20. Dijkman PM, Muñoz-García JC, Lavington SR, Kumagai PS, dos Reis RI, Yin D, et al. Conformational dynamics of a G protein-coupled receptor helix 8 in lipid membranes. Sci Adv. 2020: 6: eaav8207.

21. Proudfoot AEI. Chemokine receptors: multifaceted therapeutic targets. Nat Rev Immunol. 2002;2: 106–115.

22. Rosenkilde MM, Schwartz TW. The chemokine system -a major regulator of angiogenesis in health and disease. APMIS 2004;112: 481–495.

23. Rajagopalan L, Rajarathnam K. Structural basis of chemokine receptor function -a model for binding affinity and ligand selectivity. Biosci Rep. 2006;26: 325–339.

24. Monteclaro FS, Charo IF. The amino-terminal extracellular domain of the MCP-1 receptor, but not the RANTES/MIP-1α receptor, confers chemokine selectivity. evidence for a two-step mechanism for MCP-1 receptor activation. J Biol Chem. 1996;271: 19084–19092.

25. Kufareva I, Salanga CL, Handel TM. Chemokine and chemokine receptor structure and interactions: implications for therapeutic strategies. Immunol Cell Biol. 2015;93: 372383.

26. Rajarathnam K, Sykes BD, Kay CM, Dewald B, Geiser T, Baggiolini M, et al. Neutrophil activation by monomeric interleukin-8. Science. 1994;264: 90–92.

27. Skelton NJ, Quan C, Reilly D, Lowman H. Structure of a CXC chemokine-receptor fragment in complex with interleukin-8. Structure 1999;7: 157–168.

28. Berkamp S, Park SH, De Angelis AA, Marassi FM, Opella SJ. Structure of monomeric interleukin-8 and its interactions with the N-terminal binding site-I of CXCR1 by solution NMR spectroscopy. J Biomol NMR. 2017;69: 111–121.

29. Kufareva I, Gustavsson M, Zheng Y, Stephens BS, Handel TM. What do structures tell us about chemokine receptor function and antagonism? Annu Rev Biophys. 2017;46: 175–198.

30. Arimont M, Sun S-L, Leurs R, Smit M, de Esch IJP, de Graaf C. Structural analysis of chemokine receptor-ligand interactions. J Med Chem. 2017;60: 4735–4779.

31. Arimont M, Hoffmann C, de Graaf C, Leurs R. Chemokine receptor crystal structures: what can be learned from them? Mol Pharmacol. 2019;96: 765–777.

32. Kleist AB, Getschman AE, Ziarek JJ, Nevins AM, Gauthier P-A, Chevigné A, et al. New paradigms in chemokine receptor signal transduction: moving beyond the two-site model. Biochem Pharmacol. 2016;114: 53–68.

33. Baggiolini M, Dewald B, Moser B. Interleukin-8 and related chemotactic cytokines-CXC and CC chemokines. Adv Immunol. 1994;55: 97–179.

34. Park SH, Das, BB, Casagrande F, Tian Y, Nothnagel HJ, Chu M, et al. Structure of the chemokine receptor CXCR1 in phospholipid bilayers. Nature. 2012;491: 779–783.

35. Rajarathnam K, Clark-Lewis I, Sykes BD. ^1^H NMR solution structure of an active monomeric interleukin-8. Biochemistry. 1995;34: 12983–12990.

36. Rajagopalan L, Rajarathnam K. Ligand selectivity and affinity of chemokine receptor CXCR1. Role of N-terminal domain. J Biol Chem. 2004;279: 30000–30008.

37. Park SH, Casagrande F, Cho L, Albrecht L, Opella SJ. Interactions of interleukin-8 with the human chemokine receptor CXCR1 in phospholipid bilayers by NMR spectroscopy. J Mol Biol. 2011;414: 194–203.

38. Haldar S, Raghuraman H, Namani T, Rajarathnam K, Chattopadhyay A. Membrane interaction of the N-terminal domain of chemokine receptor CXCR1. Biochim Biophys Acta Biomembr. 2010;1798: 1056–1061.

39. Chaudhuri A, Basu P, Haldar S, Kombrabail M, Krishnamoorthy G, Rajarathnam K, et al. Organization and dynamics of the N-terminal domain of chemokine receptor CXCR1 in reverse micelles: effect of graded hydration. J Phys Chem B 2013;117: 1225–1233.

40. Kharche S, Joshi M, Sengupta D, Chattopadhyay A. Membrane-induced organization and dynamics of the N-terminal domain of chemokine receptor CXCR1: insights from atomistic simulations. Chem Phys Lipids. 2018;210: 142–148.

41. Attwood MR, Borkakoti N, Bottomley GA, Conway EA, Cowan I, Fallowfield AG, et al. Identification and characterisation of an inhibitor of interleukin-8: a receptor based approach. Bioorg Med Chem Lett. 1996;6: 1869–1874.

42. Joseph PRB, Spyracopoulos L, Rajarathnam K. Dynamics-derived insights into complex formation between the CXCL8 monomer and CXCR1 N-terminal domain: an NMR study. Molecules. 2018;23: 2825.

43. Hébert CA, Vitangcol RV, Baker JB. Scanning mutagenesis of interleukin-8 identifies a cluster of residues required for receptor binding. J Biol Chem. 1991;266: 18989–18994.

44. Hammond MEW, Shyamala V, Siani MA, Gallegos CA, Feucht PH, Abbott J, et al. Receptor recognition and specificity of interleukin-8 is determined by residues that cluster near a surface-accessible hydrophobic pocket. J Biol Chem. 1996;271: 8228–8235.

45. Schraufstätter IU, Ma M, Oades ZG, Barritt DS, Cochrane CG. The role of Tyr^13^ and Lys^15^ of interleukin-8 in the high affinity interaction with the interleukin-8 receptor type A. J Biol Chem. 1995;270: 10428–10431.

46. Williams G, Borkakoti N, Bottomley GA, Cowan I, Fallowfield AG, Jones PS, et al. Mutagenesis studies of interleukin-8. identification of a second epitope involved in receptor binding. J Biol Chem. 1996;271: 9579–9586.

47. Suetomi K, Lu Z, Heck T, Wood TG, Prusak DJ, Dunn KJ, et al. Differential mechanisms of recognition and activation of interleukin-8 receptor subtypes. J Biol Chem. 1999;274: 11768–11772.

48. Kharche SA, Sengupta D. Dynamic protein interfaces and conformational landscapes of membrane protein complexes. Curr Opin Struct Biol. 2020;61: 191–197.

49. Liu K, Wu L, Yuan S, Wu M, Xu Y, Sun Q, et al. Structural basis of CXC chemokine receptor 2 activation and signalling. Nature 2020;585: 135–140.

50. Gustavsson M, Dyer DP, Zhao C, Handel TM. Kinetics of CXCL12 binding to atypical chemokine receptor 3 reveal a role for the receptor N terminus in chemokine binding. Sci Signal. 2019;12: eaaw3657.

51. Ziarek JJ, Volkman BF. NMR in the analysis of functional chemokine interactions and drug discovery. Drug Discov Today Technol. 2012;9: e293–e299.

52. Liou J-W, Chang F-T, Chung Y, Chen W-Y, Fischer WB, Hsu H-J. *In silico* analysis reveals sequential interactions and protein conformational changes during the binding of chemokine CXCL-8 to its receptor CXCR1. PLoS One. 2014;9: e94178.

53. Helmer D, Rink I, Dalton JAR, Brahm K, Jöst M, Nargang TM et al. Rational design of a peptide capture agent for CXCL8 based on a model of the CXCL8:CXCR1 complex. RSC Adv. 2015;5: 25657–25668.

54. Jiang S-J, Liou J-W, Chang C-C, Chung Y, Lin L-F, Hsu H-J. Peptides derived from CXCL8 based on *in silico* analysis inhibit CXCL8 interactions with its receptor CXCR1. Sci Rep. 2016;5: 18638.

55. Delort B, Renault P, Charlier L, Raussin F, Martinez J, Floquet N. Coarse-grained prediction of peptide binding to G-protein coupled receptors. J Chem Inf Model. 2017;57: 562–571.

56. Lelimousin M, Limongelli V, Sansom MSP. Conformational changes in the epidermal growth factor receptor: role of the transmembrane domain investigated by coarse-grained metadynamics free energy calculations. J Am Chem Soc. 2016;138: 10611–10622.

57. Pawar AB, Sengupta D. Resolving the conformational dynamics of ErbB growth factor receptor dimers. J Struct Biol. 2019;207: 225–233.

58. Prasanna X, Chattopadhyay A, Sengupta D. Cholesterol modulates the dimer interface of the β_2_-adrenergic receptor via cholesterol occupancy sites. Biophys J. 2014;106: 1290–1300.

59. Prasanna X, Sengupta D, Chattopadhyay A. Cholesterol-dependent conformational plasticity in GPCR dimers. Sci Rep. 2016;6: 31858.

60. Sengupta D, Prasanna X, Mohole M, Chattopadhyay A. Exploring GPCR-lipid interactions by molecular dynamics simulations: excitements, challenges, and the way forward. J Phys Chem B. 2018;122: 5727–5737.

61. Prasanna X, Mohole M, Chattopadhyay A, Sengupta D. Role of cholesterol-mediated effects in GPCR heterodimers. Chem Phys Lipids. 2020;227: 104852.

62. Marrink SJ, Risselada HJ, Yefimov S, Tieleman DP, de Vries AH. The MARTINI force field: coarse grained model for biomolecular simulations. J Phys Chem B. 2007;111: 7812–7824.

63. de Jong DH, Singh G, Bennett WFD, Arnarez C, Wassenaar TA, Schäfer LV, et al. Improved parameters for the Martini coarse-grained protein force field. J Chem Theory Comput. 2013;9: 687–697.

64. Wassenaar TA, Ingólfsson HI, Böckmann RA, Tieleman DP, Marrink SJ. Computational lipidomics with *insane:* a versatile tool for generating custom membranes for molecular simulations. J Chem Theory Comput. 2015:11: 2144–2155.

65. Periole X, Cavalli M, Marrink S-J, Ceruso MA. Combining an elastic network with a coarse-grained molecular force field: structure, dynamics, and intermolecular recognition. J Chem Theory Comput. 2009;5: 2531–2543.

66. Van Der Spoel D, Lindahl E, Hess B, Groenhof G, Mark AE, Berendsen HJC. GROMACS: fast, flexible, and free. J Comput Chem. 2005;26: 1701–1718.

67. Pronk S, Páll S, Schulz R, Larsson P, Bjelkmar P, Apostolov R, et al. GROMACS 4.5: a high-throughput and highly parallel open source molecular simulation toolkit. Bioinformatics. 2013;29: 845–854.

68. Bussi G, Donadio D, Parrinello M. Canonical sampling through velocity rescaling. J Chem Phys. 2007;126: 014101.

69. Berendsen HJC, Postma JPM, van Gunsteren WF, DiNola A, Haak JR. Molecular dynamics with coupling to an external bath. J Chem Phys. 1984;81: 3684.

70. Wu EL, Cheng X, Jo S, Rui H, Song KC, Dávila-Contreras EM, et al. CHARMM-GUI *membrane builder* toward realistic biological membrane simulations. J Comput Chem. 2014;35: 1997–2004.

71. Parrinello M, Rahman A. Polymorphic transitions in single crystals: a new molecular dynamics method. J Appl Phys. 1981;52: 7182.

72. Humphrey W, Dalke A, Schulten K. VMD: visual molecular dynamics. J Mol Graph. 1996;14: 33–38.

73. Han B, Liu Y, Ginzinger SW, Wishart DS. SHIFTX2: significantly improved protein chemical shift prediction. J Biomol NMR. 2011;50: 43–57.

